# Separation-of-function mutants reveal the NF-κB-independent involvement of IκBα in the regulation of stem cell and oncogenic programs

**DOI:** 10.1101/2023.06.21.545928

**Authors:** Daniel Álvarez-Villanueva, Luis Galán-Palma, Joan Bertran, Martin Floor, Laura Solé, Teresa Lobo-Jarne, María Maqueda, Rajani Rajbhandari, Laura Marruecos, Jordi Villà-Freixa, Markus Bredel, Anna Bigas, Lluís Espinosa

## Abstract

We previously demonstrated that the NF-κB inhibitor IκBα binds the chromatin together with PRC2 to regulate a subset of developmental- and stem cell-related genes. This alternative function has been elusive in both physiological and disease conditions because of the predominant role of IκBα as a negative regulator of NF-κB.

We here uniquely characterize specific residues of IκBα that allow the generation of separation-of-function (SOF) mutants that are defective for either NF-κB-related (SOF^ΔNF-κB^) or chromatin-related (SOF^ΔH2A,H4^) activities. Expression of IκBα SOF^ΔNF-κB^, but not SOF^ΔH2A/H4^, is sufficient to negatively regulate a specific stemness program in intestinal cells, thus rescuing the differentiation blockage imposed by IκBα deficiency. In contrast, full IκBα activity is required for regulating clonogenic/tumor-initiating activity of colorectal cancer cells.

Our data indicate that SOF mutants represent an exclusive tool for studying IκBα functions in physiology and disease, and identified IκBα as a robust prognosis biomarker for human cancer.

## Introduction

The NF-κB pathway is an essential regulator of inflammation, immune response and cellular survival, being the IκB family of proteins the primary inhibitors of this pathway both under non-stimulated conditions and following activation^1^. Canonical NF-κB, initiated by stimulus such as TNFα, leads to activation of the IKK kinase complex that phosphorylates IκBs at specific serine (S) residues (S32 and S36 in the case of IκBα), thus inducing ß-TRCP-dependent K48-linked ubiquitination and proteasomal degradation. For years, negative regulation of NF-κB has been considered the only function for IκBα. However, alternative post-translational modifications of IκBα have also been shown to affect its activity under certain conditions. SUMOylation of IκBα by SUMO1 was initially recognized to prevent its TNFα-induced degradation and thus NF-κB activation^2^. Contrarily, SUMO2/3 linkage to IκBα induced by hypoxia facilitates p65 release leading to NF-κB activation^3^. Other stimuli such as H_2_O_2_, epidermal growth factor or pervanadate induce tyrosine (Y) phosphorylation of IκBα (at Y42), leading to NF-κB activation independent of IκBα degradation^4–6^. Phosphorylation of IκBα at Y42 is required for proper neuronal development^7^. Alternative non-degradative phosphorylation of Y289 and Y305 of IκBα promotes NF-κB activation in B cells^8^.

Previous results from our group demonstrated that SUMOylated IκBα is predominantly nuclear and associates with histones in the chromatin to modulate the activity of polycomb repressor complex 2 (PRC2) at a subset of stemness- and developmental-related genes. Association of IκBα with NF-κB or histone H4 *in vitro* is mutually exclusive^9^, suggesting that a common domain in IκBα mediates both interactions. Further investigations indicated that developmental defects associated with IκBα deficiency are linked to its chromatin and PRC2-related function^9–1^^1^. Due to the predominant role of IκBα as regulator of NF-κB, the broad spectrum of NF-κB activities and the multiple crosstalk between NF-κB and PRC2 activities^12–15^, uncovering the functional impact of chromatin-related IκBα in specific systems has remained extremely challenging. This is the case of human cancers such as glioblastoma^16, 17^, low-grade glioma^18^, non-small cell lung carcinoma^19^, gray zone lymphoma^20^ or Hodgkin lymphoma^21–23^, and different murine models of skin cancer^9, 24^, in which IκBα deficiency has recurrently been observed. Revealing the relative impact of NF-κB- or PRC2-related IκBα functions in these systems will have relevant clinical implications.

Here, we have defined a computational metric, the fold-excluded evolutionary conservation (FEEC), which combines domain compactness (through the weighted contact number -WCN) with residue conservation to identify the residues in the IκBα protein that define its association with NF-κB and histones. We have then generated separation-of-function mutants and used them to demonstrate the differential requirement of chromatin- and NF-κB-related IκBα functions for transcriptional regulation, intestinal cell differentiation and tumor initiating capacity of colorectal cancer cells.

## Results

### A common domain of IκBα binds to p65 and histone H4

We aimed to investigate the determinants of IκBα association with NF-κB or chromatin/histones. Based on the strict evolutionary conservation of histone tails, we hypothesized that residues of IκBα involved in histone binding would also be similarly conserved. To differentiate IκBα residues that are conserved because they participate in other functions besides protein folding (e.g., in protein-protein binding, for surface exposed residues), we derived a classification score named FEEC (fold-<u>e</u>xcluded <u>e</u>volutionary <u>c</u>onservation) that allows identifying conserved residues with functional roles beyond maintaining the structural fold of the protein (see Methods for the FEEC score definition). From the set of 213 IκBα residues in the crystallographic structure (PDB code: 1NFI, encompassing residues 70 to 282 in the IκBα sequence; Uniprot entry P25963), we considered 132 residues to have positive FEEC values (Figure 1A, upper panels and Table S1), of which around half (65) are solvent-exposed —solvent-accessible surface area (SASA) higher than 0.2 nm2— and could putatively participate in protein-protein interactions.

**Figure 1.**
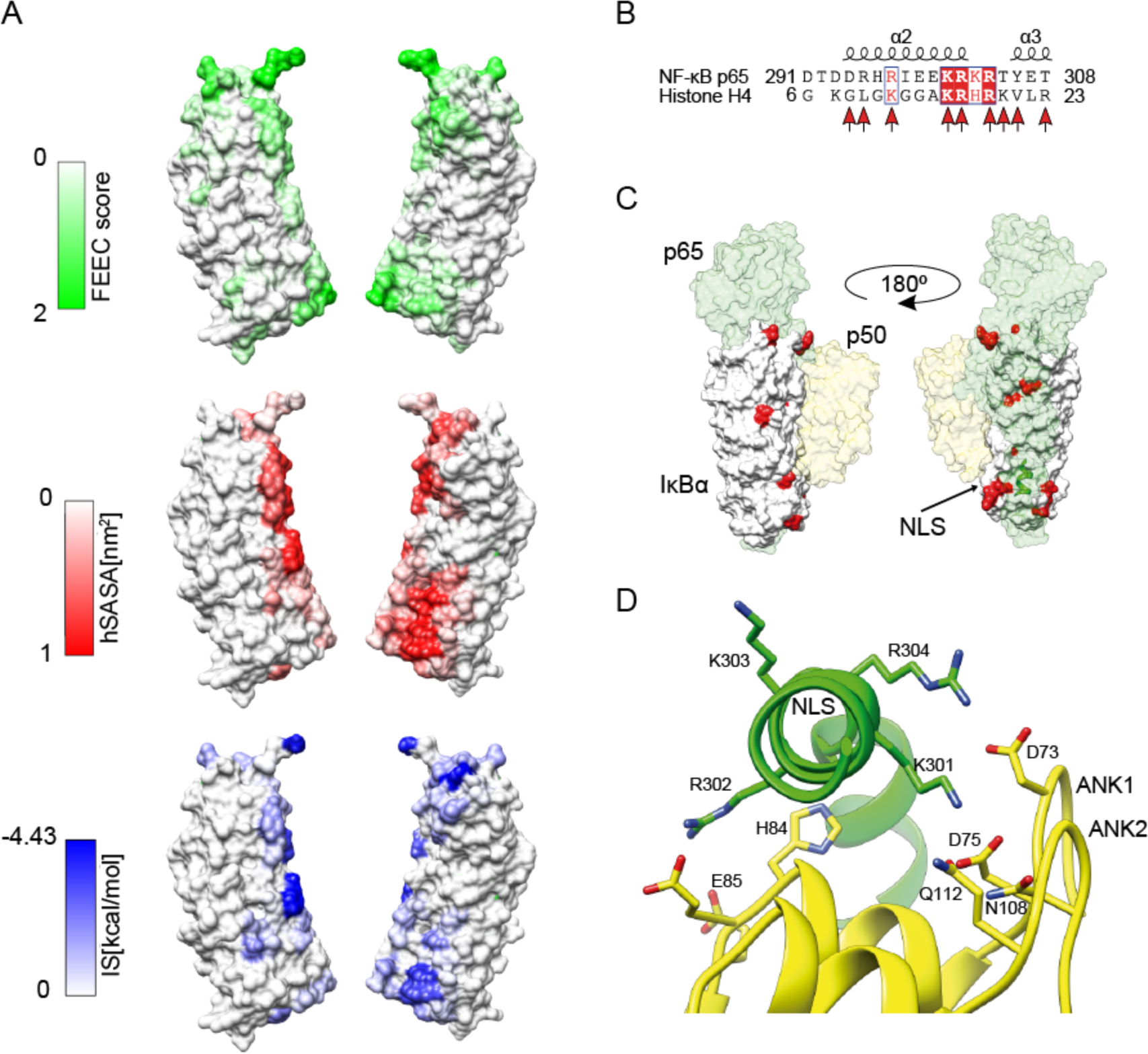
A common domain of IκBα is required for p65 and histone H4 binding. **(A)** (Upper) Surface mapping of residues with positive fold-excluded evolutionary conservation scores (maximum value capped at 2 to exclude outliers). (Middle) Fraction of the solvent-accessible surface area hidden upon complexation of the IκBα and NF-κB complex structure. (Bottom) Per-residue interface scores of the IκBα and NF-κB complex structure. The structures are related by a 180° turn— PDB structure 1NFI. **(B)** Alignment of the NF-κB/p65 NLS region to H4 N-terminal tail. Similar positions are indicated in blue rectangles; similar and identical residues, inside a position, are in red and white letters respectively. Red arrows indicate NF-κB/p65 residues participating in IκBα binding. Secondary structure is from PDB structure 3UW9. **(C)** IκBα (white) and NF-κB/p65 (green) and p50 (yellow) complex. All negatively charged IκBα residues with a positive FEEC score appear as red surfaces. The NF-κB/p65 Nuclear Localization Signal (NLS) region is indicated. **(D)** Interaction of NF-κB/p65 NLS motif with IκBα ANK1 and ANK2 repeats. Polar IκBα residues interacting with the NLS are depicted in yellow, NLS motif, KRKR, is shown in green. They are numbered according to Uniprot entries Q04206 (NF-κB/p65) and P25963 (IκBα) — PDB structure 1NFI.

To validate our model, we classified which residues in the IκBα/NF-κB interface could be predicted using the FEEC score alone —interface residues were defined as having more than 20% of their SASA hidden (SASAh) upon complexation. For the interaction with NF-κB, 83% (30/36) and 73% (8/11) of IκBα interface residues interacting with the p65 and p50 subunits, respectively, were correctly included in the set of positive FEEC residues. While SASAh is derived from a geometric definition of residues belonging to the interface, it does not account for chemical interactions or their strength; residues not participating in significant interactions are still expected to be included in the interface just because nearby interface residues occlude them. Therefore, we repeated our interface analysis considering per-residue interface scores (for details, see Methods) to define their role in the IκBα and NF-κB interface more quantitatively. Our method shows that all strongly interacting residues were correctly included in our set of positive FEEC residues (Figure S1A). We then mapped the positive FEEC scores to the protein surface to compare their distribution with the location of the residues of the IκBα/NF-κB interfaces (obtained from the crystal structure of this complex^25^), defined either by the per-residue SASAh, or the interface binding score values. Interestingly, there is a strikingly similar distribution pattern between surface residues with positive FEEC scores and residues participating at the interface (Figure 1A), validating the FEEC metric as a predictor of putative binding sites for the IκBα protein.

The N-terminal tail of histone H4 includes many positively charged residues. Thus, logical surface positions for its binding should include charge complementarity (i.e., negatively charged residues). To identify hot spots for interaction with H4, we considered all the negatively charged surface IκBα residues having positive FEEC scores (Table S1). Importantly, sequence alignment between the p65-NF-κB <u>N</u>uclear <u>L</u>ocalization <u>S</u>ignal (NLS) and the H4 N-terminal tail revealed an alpha-helical motif conserved between the two proteins (Figure 1B), with different secondary structure prediction methods showing a comparable helical character for the region between K16 and K20 of histone H4 (Figure S1B). Based on these data, the most likely IκBα region for binding to histone H4 is coincident with the one binding the p65-NF-κB NLS (Figure 1C). Inside this region, NLS of p65-NF-κB makes contact with many IκBα charged and polar residues with positive FEEC scores, including D73, D75, H84, E85, E86, N108 and Q112 (Table S1) that belong to the ANK1 and ANK2 subdomains (Figure 1D). We propose that the same IκBα interface patch that interacts with the p65-NF-κB NLS could mediate the interactions with histone H4.

### Two different aminoacidic clusters in IκBα define NF-kB and chromatin interaction

Consistent with their putative functional relevance, residues D73, D75, H84, E85, E86, N108 and Q112 of IκBα (corresponding to the human sequence, Uniprot entry P25963) are highly conserved from nematodes to humans (Figure S2A) and among different IκB homologues (Figure S2B). To investigate the contribution of these residues to NF-κB and histone binding, we first carried out an in-silico mutagenesis to determine the effect of changing these positions to non-synonymous residues over the structure of IκBα. According to a mutation penalty matrix, all positions will accept an alanine substitution without significantly changing the energy of the system (Figure 2A). Therefore, we generated different compose mutants carrying alanine substitutions of residues D73, D75, H84, E85, E86, N108 and Q112 (Figure 2B) and tested them functionally in co-precipitation (Co-IP) and pull-down (PD) assays.

**Figure 2.**
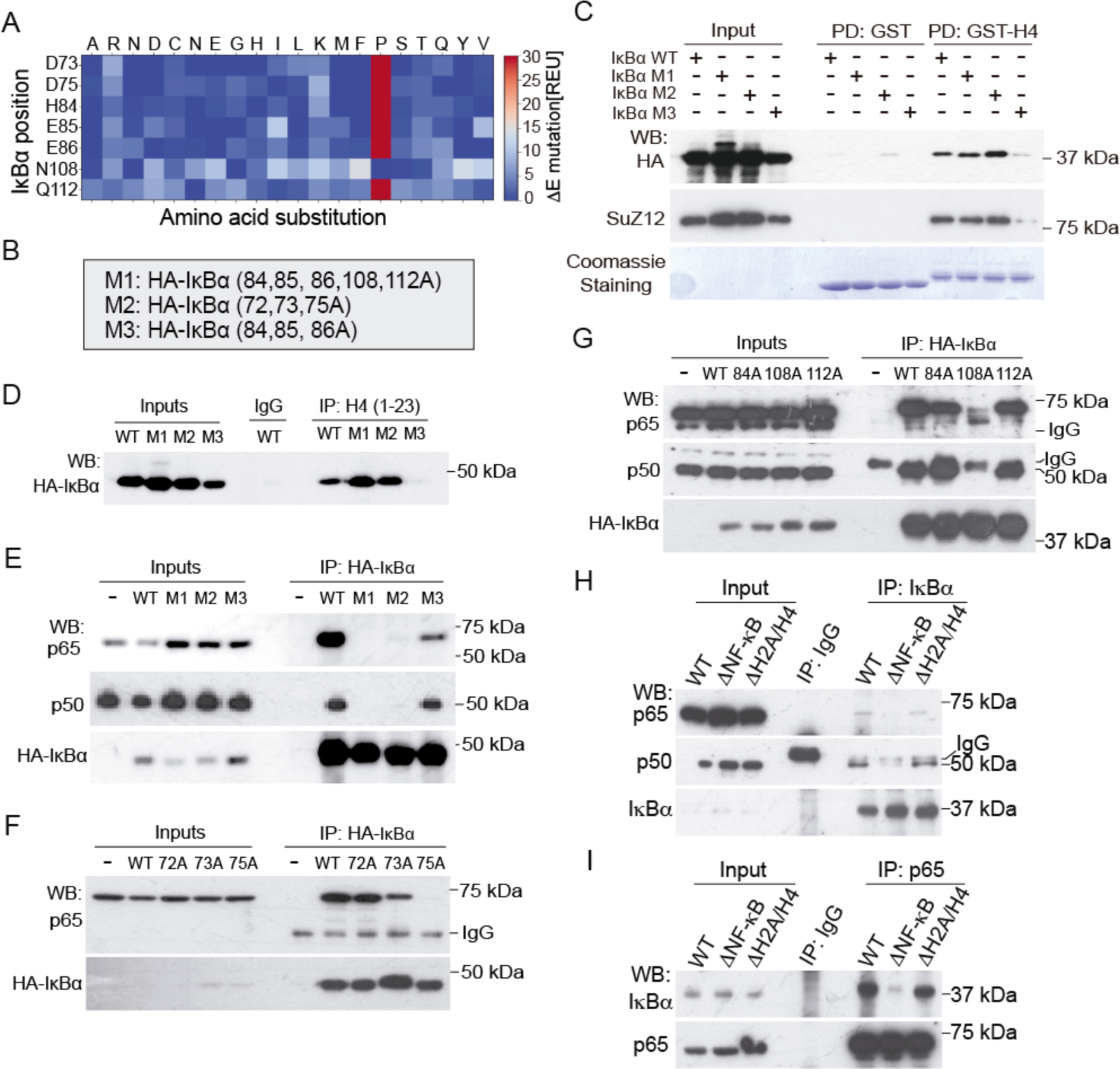
Two different aminoacidic clusters in IκBα define NF-κB and histone H4 interaction. **(A)** Analysis of the energy differences produced by mutating the WT amino acid in the indicated position to any of the other 20 amino acids. **(B)** Compose mutants generated for evaluation of binding affinity with NF-κB and histone H4. **(C, D)** Precipitation assays using GST, GST-H4 as bait (C) or biotinylated peptides corresponding to amino acids 1-23 of H4 (D) and the indicated compose IκBα mutants expressed in HCT116 or HEK-293T cells. We determined the presence of IκBα in the precipitates by WB with anti-HA antibody. Coomassie staining in C shows comparable amount of GST or GST-H4 in the assays. **(E)** WB analysis of the indicated NF-κB subunits in the precipitates of the indicated HA-IκBα mutants expressed in HCT116 cells. **(F, G)** PD (F) and Co-IP (G) analysis of the indicated IκBα single point mutants. Notice the total abrogation of p65 and p50 association with the single N108A and D75A mutants in HCT116 cells. **(H, I)** WB analysis of the indicated proteins in the precipitates of IκBα (H) and NF-κB/p65 (I) from HCT116 IκBα knock-in lysates.

By PD assay of HEK-293T cell lysates, we found that mutating N72, D73, and D75 to alanine (M2 mutant) did not affect IκBα binding to histones H4 and H2A, which was almost abolished by the H84, E85, E86 to alanine mutations (Figures 2C and S2C). Notably, defective association of M3 to histones, which is imposed by H84, E85 and E86 to A mutations, was reverted by the additional substitution of N108 and Q112 to A (see M1 in Figure 2C), suggesting that N108 or Q112 positions may favor the competing NF-κB association. Identical results were obtained by co-IP with a peptide corresponding to the N terminal tail of histone H4 (aa 1-23) (Figure 2D). We performed Co-IP assays from transfected HEK-293T cells to test the binding capacity of IκBα mutants with endogenous NF-κB subunits. We found reduced association of the M3 mutant to p65/NF-κB compared with wildtype (WT) IκBα, whereas mutants M1 (N72A, D73A, D75A) and M2 (H84A, E85A, E86A, N108A, Q112A) displayed an absolute lack of p65 and p50 NF-κB binding (Figure 2E). In the same set of experiments, we demonstrated that only histone binding-proficient IκBα proteins increased SUZ12 association to histone H4 (Figure 2C), as we previously shown for WT IκBα^9^, further supporting the concept that chromatin-bound IκBα regulates PRC2 function.

We then generated single-point mutants for residues candidate to impose IκBα binding specificity. By PD and Co-IP experiments, we found that D75A or N108A mutations were sufficient to disrupt IκBα association to NF-κB specifically (Figure 2F). In contrast, single mutations of H84, E85 or E86 to alanine did not affect histone H4 binding compared with the triple H84A, E85A and E86A mutant (Figure 2G). We then mutated the IκBα protein to introduce the changes present in IκBβ in the (HEE to HQH) and tested the hybrid molecule (from now IκBα/β-mimic) in IP and PD assays. IκBα/β-mimic bound to p65 comparable to IκBα (Figure S2D), as expected, but show a significant defective association to histone H4 (Figure S2E). In the same pull-down assay, we did not detect any association of endogenous IκBε (that show a single E46 to A substitution compared with IκBα, see Figure S2B) to histone H4 (Figure S2D). Since, E46 to A substitution in the IκBα protein did not preclude histone binding, this result indicate that additional residues contribute to regulate IκB to histone interaction. From now on, we will call IκBα SOF^ΔNF-κB^ those mutants displaying defective NF-κB binding (but retaining histone binding) (M1, M2, D75A and N108A) and SOF^ΔH2A/H4^ the IκBα mutant that fails to bind histones but associates with p65 and p50 (M3 and IκBα/β-mimic). To exclude the possible impact of protein over-expression in the interaction results, we generated IκBα knock-in HCT116 CRC cells carrying the N108A (SOF^ΔNF-κB^) or HEE to HQH (IκBα/β-mimic=SOF^ΔH2A/H4^) mutations. We found that the SOF^ΔNF-κB^ mutant expressed under the control of the endogenous *NFKBIA* promoter failed to interact with p65 and p50, similar to the ectopically expressed protein, and in contrast with IκBα WT and SOF^ΔH2A/H4^ (Figures 2H and 2I).

Together, these results support the concept that p65-NF-κB and histone H4 compete for binding to the same IκBα region with specific residues differentially influencing its association with p65 -NF-κB and histones.

### IκBα mutants display separation of function (SOF) activity in cells

To study IκBα SOF mutants functionally, we reconstituted IκBα knockout (KO) HT-29 M6 cells (CRC cells) with doxycycline-inducible (i) IκBα WT, IκBα SOF^ΔNF-κB^ (we used N108A) and SOF^ΔH2A/H4^ (H84A, E85A, E86A) constructs. We first tested the capacity of i-IκBα WT or SOF mutants (induced by 16 hours of doxycycline treatment) (Figure S3) to prevent nuclear translocation of p65/RelA upon TNFα treatment. As determined by immunofluorescence (IF) analysis, ectopic and sustained expression of i-IκBα WT completely abrogated nuclear p65 translocation after 30 and 60 minutes of TNFα treatment (Figure 3A and 3B). A comparable but partial effect was obtained upon expression of i-IκBα SOF^ΔH2A/H4^ (Figure 3B). In contrast, high levels of i-IκBα SOF^ΔNF-κB^ did not prevent p65 translocation (Figures 3A and 3B), consistent with their defective association to NF-κB factors in the *in vitro* assays. The differential effect of i-IκBα mutants on p65 chromatin recruitment upon TNFα treatment was confirmed by subcellular fractionation followed by WB analysis of HT-29 M6 cells (Figure 3C). Next, we tested the impact of ectopic i-IκBα WT or SOF mutants in the TNFα-induced transcription of the canonical NF-κB target *A20* (Figure 3D). Consistent with their negative impact over p65 nuclear translocation, both IκBα WT and the SOF^ΔH2A/H4^ mutant almost precluded transcriptional activation of *A20* after 30 minutes of TNFα treatment and reduced *A20* levels at 60 minutes when compared with IκBα SOF^ΔNF-κB^ (Figure 3D). These results indicate that IκBα SOF^ΔNF-κB^ is deficient in NF-κB regulation whereas the SOF^ΔH2A/H4^ mutant represses NF-κB activation by TNFα comparable to the WT protein.

**Figure 3.**
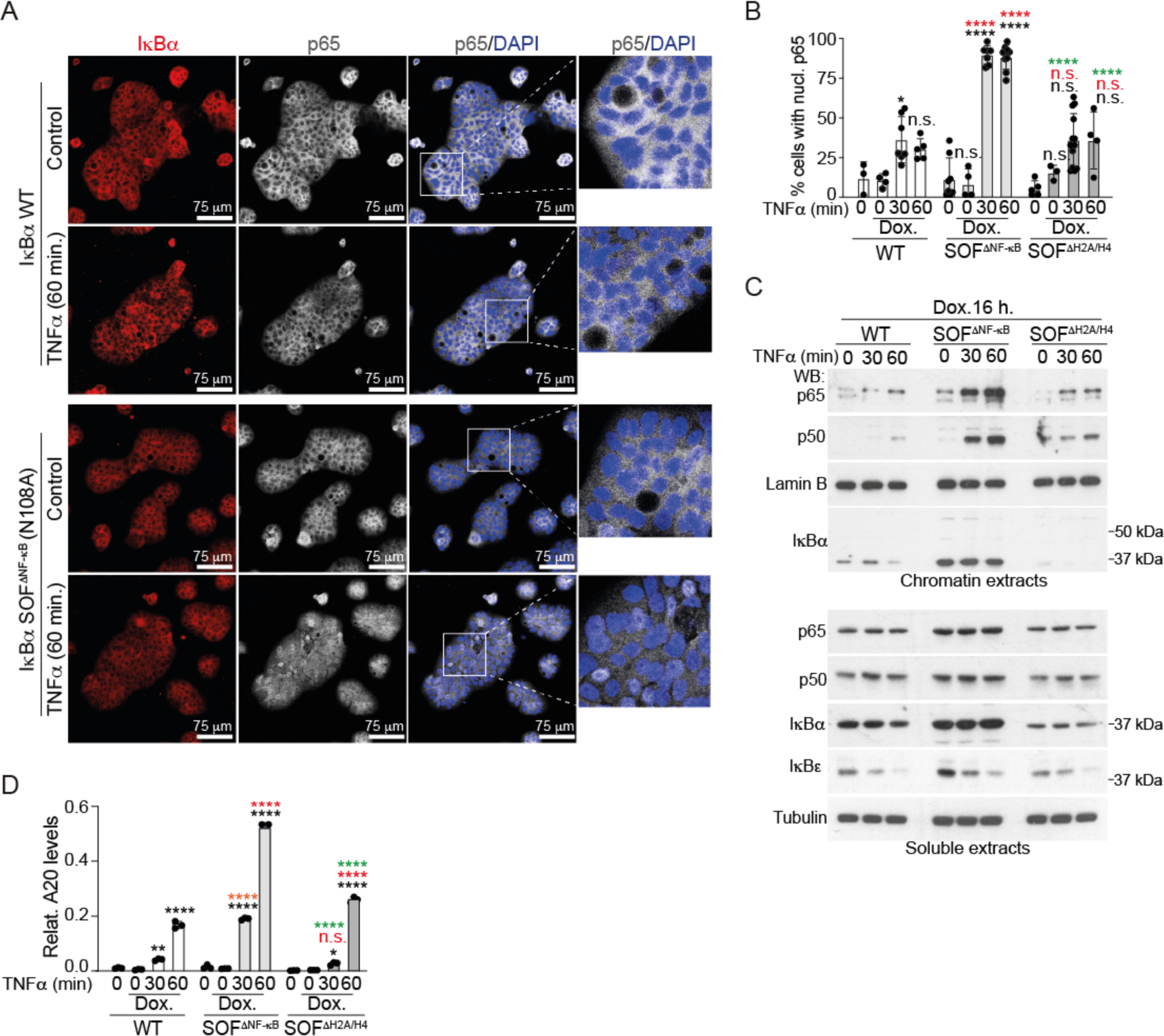
IκBα mutants display separation of function (SOF) activity in intestinal cells. **(A)** IF analysis of p65/NF-κB of IκBα-deficient HT-29 M6 cells reconstituted (by 16 hours doxycycline treatment) with the SOF^ΔNF-κB^ mutant N108A or WT IκBα proteins and then treated with TNFα as indicated. **(B)** Quantification of the percent of cells displaying nuclear p65 in a minimum of 5 colonies per condition counted. **(C)** Western blot analysis of cytoplasmic and chromatin fractions from i-IκBα WT and SOF reconstituted HT-29 M6 cells. **(D)** qPCR analysis of the canonical NF-κB target gene *A20* in IκBα KO HT-29 M6 cells reconstituted as indicated. Expression levels of the different genes were calculated relative to *GAPDH*, and statistical analysis of differences determined by Two-way Anova test comparing treatments with no-treatment (black labels), same treatments in WT (red labels) or same treatments in SOF^ΔNF-κB^ (green labels). ****p-value < 0.0001, ***p-value < 0.0005, **p-value <0.0025 *p-value < 0.05; n.s.: no significant.

### IκBα SOF^ΔNF-κB^ mutant N108A rescues the Goblet Cell differentiation blockage imposed by IκBα deletion

HT-29 M6 CRC cells acquire a goblet-cell phenotype upon confluence^26^, which is preclude by IκBα depletion, as determined by reduced MUC5AC expression at 7 days post-confluence^27^ (Figure 4A). We tested whether i-IκBα WT, SOF^ΔNF-κB^ and/or SOF^ΔH2A/H4^ recued the differentiation capacity of IκBα KO HT-29 M6 cells. We initially tested two different IκBα induction protocols: i) 16 hours of dox. treatment at pre-confluence and culture in fresh medium up to 7 dpc and ii) dox. treatment for the whole period of the experiment. We found that 16 hours of IκBα SOF^ΔNF-κB^ induction were sufficient to restore MUC5AC levels in IκBα KO HT-29 M6 cells at levels comparable to i-IκBα WT. However, SOF^ΔH2A/H4^ did not revert the defective differentiation of IκBα KO cells (Figure 4B). To uncover the molecular bases by which IκBα mutant differentially impact goblet cell differentiation capacity, we performed RNA-seq analysis of HT-29 M6 cells reconstituted with i-IκBα WT, SOF^ΔNF-κB^ and SOF^ΔH2A/H4^ for 16 hours (Figures 4C and Table S2) (GSE206515). Although we detected a significant clustering by cell lines in the Principal Component Analysis (PCA), SOF^ΔNF-κB^ showed the best separation when comparing doxycycline-treated and untreated cells, followed by IκBα WT (Figure 4D). Further evaluation of differentially expressed genes (DEG) following doxycycline induction demonstrated that SOF^ΔNF-κB^ imposed the highest transcriptional effect leading to downregulation of 652 genes and upregulation of 278 genes considering an adjusted p-value of p<0.01 (Figure 4E). We only detected 1 gene, CEMIP, differentially expressed upon ectopic expression of SOF^NF-κB^, indicating that transcriptional control by IκBα in HT-29 M6 cells is IκBα dependent but NF-κB independent. IκBα WT imposed an intermediate effect on gene transcription (Figure 4E). Importantly, Gene Set Enrichment Analysis (GSEA) of DEG from i-IκBα WT and SOF^ΔNF-κB^ reconstituted cells show relevant differences. Most notably, whereas we revealed a significant inverse correlation (p<0.05) between i-IκBα WT expression and TNFA_SIGNALING_VIA_NFKB, as expected, we did not detect any alteration associated with this pathway in the i-IκBα SOF^ΔNF-κB^ expressing cells (Figure 4F). In the same direction, analysis of CHEA dataset to search for factors likely contributing to transcriptional regulation of the top 50 repressed genes upon i-IκBα SOF^ΔNF-κB^ expression failed to detect any NF-κB-related factor. However, we identified the putative chromatin repressor PPAR delta^28^, the EZH2 associated factors TRIM28^29^, SOX2 and MYC^30, 31^, and the PRC2 subunit SUZ12 as possible factors involved in IκBα SOF^ΔNF-κB^-mediated gene repression (Figure 4G). To test whether gene repression and intestinal differentiation induced by doxycycline inducible (i)-IκBα SOF^ΔNF-κB^ expression was associated with altered H3K27me3 patterns, we performed ChIP-seq analysis of IκBα knock out cells reconstituted with i-IκBα SOF^ΔNF-κB^, untreated or treated with doxycycline for 16h (n=3 per condition). We detected 27,864 and 26,793 gene-associated regions that were H3K27 methylated in untreated and doxycycline-treated HT29 cells, respectively, corresponding to 9,290 and 9,529 annotated genes. From these genes, 8,508 were found methylated in both conditions, 782 specifically methylated in basal conditions and 1097 upon doxycycline treatment. Among the in-common methylated genes, 2,071 (24%) showed differential H3K27me3 levels in untreated and doxycycline-treated cells (considering an adjusted p-value of p<0.05). Interestingly, from the 3,650 genomic regions that were differentially methylated upon doxycycline treatment, 645 were hypomethylated and 3,005 hypermethylated indicating that expression of i-IκBα SOF^ΔNF-κB^ positively impacts on PRC2 activity. We then studied whether transcriptional regulation observed following i-IκBα SOF^ΔNF-κB^ expression was associated with changes in H3K27me3 (Table S3) (GSE206515). Counterintuitively, we found that genes differentially expressed in doxycycline-treated i-IκBα SOF^ΔNF-κB^ cells were non-randomly ascribed to the group of genes that were non detected as H3K27 methylated (Figure 4H), suggesting that transcriptional repression by i-IκBα SOF^ΔNF-κB^ initially takes place in genes with low methylation levels and it was independent, or previous, to major changes in H3K27 methylation, in agreement with our previously observations^9^. However, of 142 DEGs with H3K27me3 mark in both experimental conditions, a significant proportion showed a slight but significant increase in methylation upon i-IκBα SOF^ΔNF-κB^ expression including several intestinal stem cell genes such as *ADRA2A*, *SCL7A1* or *ASCL2* (Table S2).

**Figure 4.**
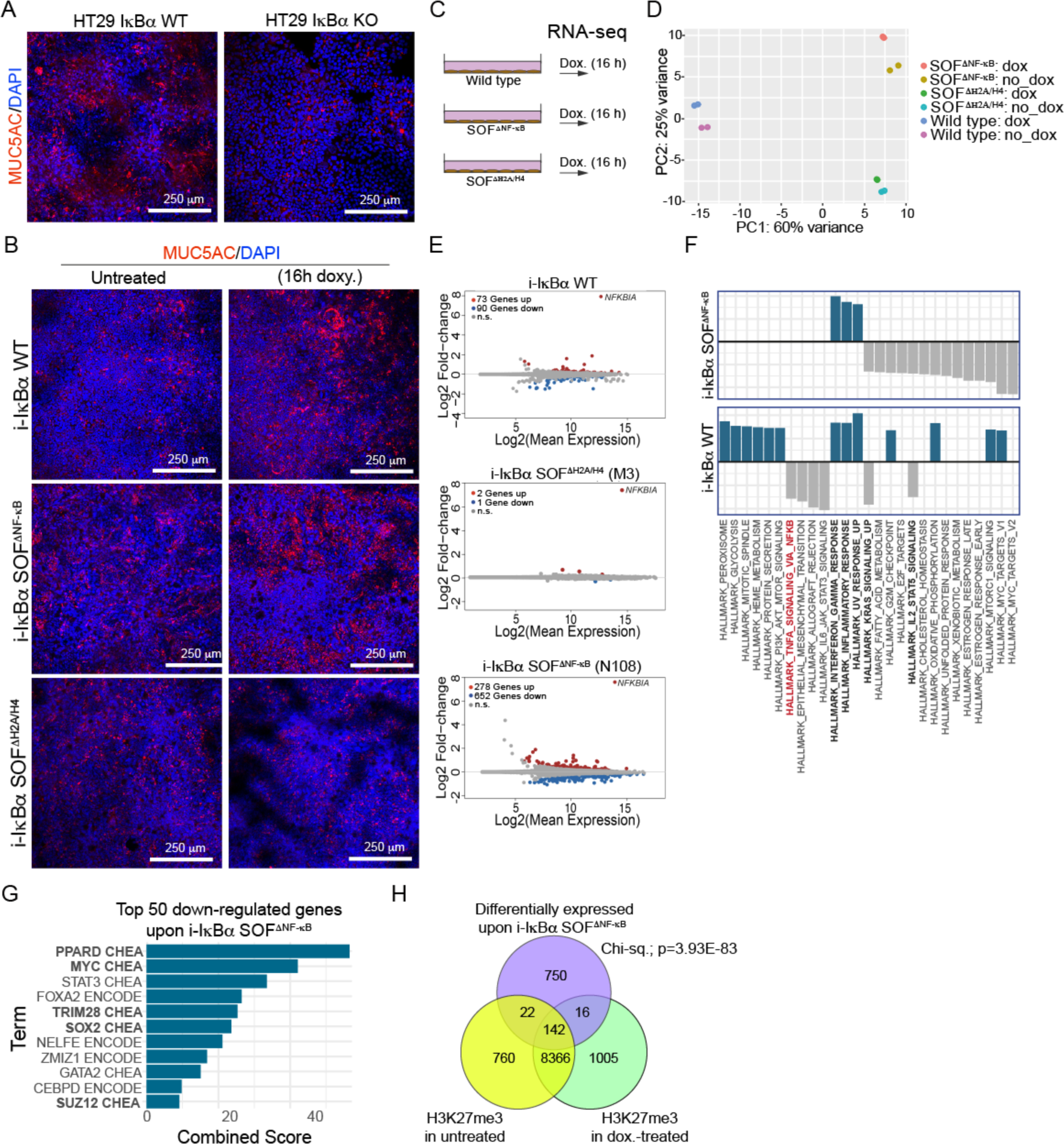
IκBα SOF^ΔNF-κB^ mutant N108A rescues the Goblet Cell differentiation blockage imposed by IκBα deletion. **(A, B)** IF analysis of WT and IκBα KO HT-29 M6 cells (A) and doxycycline-inducible IκBα WT, IκBα SOF^ΔNF-κB^ and SOF^ΔH2A/H4^ lines (B). **(C)** Scheme of the strategy used to study the transcriptional impact of i-IκBα WT and SOF expression. **(D)** Principal Component Analysis of the RNA-seq data **(E)** Graphical representation of genes differentially expressed in the different doxycycline-inducible HT-29 M6 models. **(F)** Barplot depicting the normalized enrichment score of statistically significant enriched pathways obtained by GSEA analysis with the Hallmark gene set for genes differentially repressed upon doxycycline treatment of i-IκBα SOF^ΔNF-κB^ and i-IκBα WT. **(G)** CHEA analysis of the top 50 genes that were downregulated in the doxycycline-treated IκBα SOF^ΔNF-κB^ to search for candidate transcription factors of repressor involved in IκBα-mediated gene repression. Note the absence of any NF-κB subunit in the analysis. **(H)** Venn diagram showing the overlap between genes differentially expressed upon 16 hours of doxycycline treatment of i-IκBα SOF^ΔNF-κB^ cells and genes with detectable levels of H3K27me3 in the same cells (see Table S3). Notice that most of the genes with differential expression upon i-IκBα SOF^ΔNF-κB^ correspond to genes with low methylation levels.

These results are consistent with the concept that transcriptional regulation imposed by IκBα SOF^ΔNF-κB^ was independent of NF-κB but linked to chromatin-related IκBα activities.

### Rescue of intestinal differentiation by IκBα SOF^ΔNF-κB^ is linked to repression of the ISC transcriptional program

We reasoned that the capacity of i-IκBα WT and SOF^ΔNF-κB^ to restore the differentiation capacity of HT-29 M6 cells could be associated with a negative regulation of the intestinal stem cell program. In agreement with this notion, transcriptional changes imposed by both i-IκBα WT and SOF^ΔNF-κB^ expression inversely correlated with the Intestinal Stem Cell (ISC) signature from Muñoz et al^32^ (Figure 5A and 5B); in fact, 42 out of 124 genes in this stem cell signature were significantly repressed upon IκBα SOF^ΔNF-κB^ expression including the canonical stem cell genes *MYC*, *CD44*, *CDCA7* or *PROM1*. In contrast, none of the genes upregulated upon SOF^ΔNF-κB^ expression were included in the ISC signature (Figure 5C, S5A and S5B). Downregulation of ISC genes by i-IκBα was previous to the induction of the differentiation markers MUC2 and MUC5AC (Table S2). By qPCR analysis of 16 hours doxycycline-treated WT, SOF^ΔNF-κB^ and SOF^ΔH2A/H4^ HT-29 M6 expressing cells, we confirmed that several ISC genes were repressed in an NF-κB-independent manner (SOF^ΔNF-κB^ is sufficient to impose gene repression) whereas others, such as *LGR5*, required the full IκBα activity (only repressed by IκBα WT) (Figure S5C).

**Figure 5.**
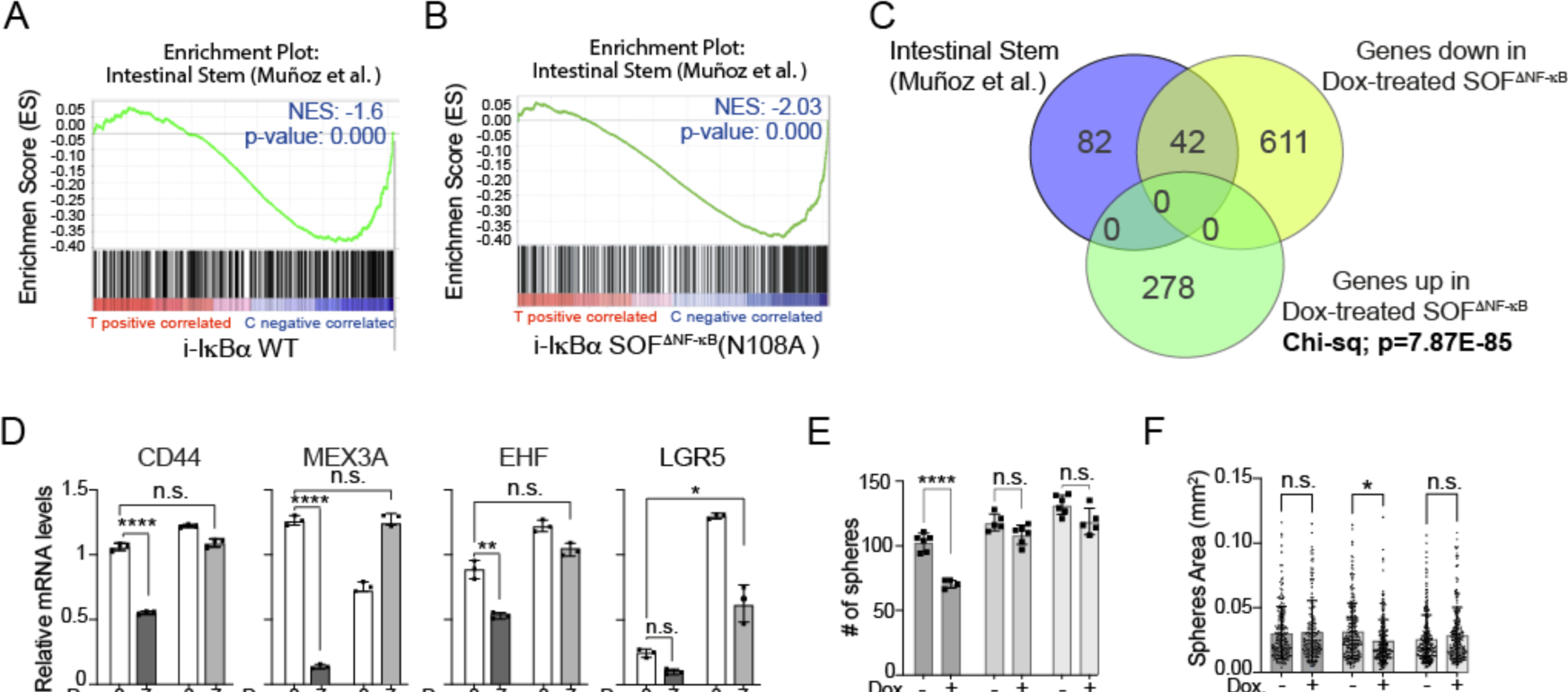
i-IκBα WT and SOF^ΔNF-κB^ expression regulates a specific intestinal stem cell program with impact on intestinal cell differentiation. **(A, B)** GSEA of an intestinal stem cell (ISC) gene set, according to Muñoz et al^32^, from genes significantly repressed upon ectopic expression (16 hours) of i-IκBα WT (A) or i-IκBα SOF^ΔNF-κB^ (B). **(C)** Venn diagram representing the overlap between genes downregulated following i-IκBα SOF^ΔNF-κB^ expression and genes in the ISC signature from Muñoz et al^32^. The significance of the correlation was determined by Chi-square analysis considering a total number of 20,500 genes in the mammalian genome^51^. **(D)** qPCR analysis of the indicated ISC genes in WT and IκBα KO HT-29 M6 cells at pre-confluence or 7 days of post-confluence (Dpc). **(E, F)** Number of organoids (E) and average organoid area (F) obtained by seeding 600 HT-29 M6 cells carrying the indicated i-IκBα constructs and left untreated or treated with doxycycline (Dox.) for 16 hours before seeding. Same results were obtained by treating cells for the whole culture period (7 days). In D-F, bars represent mean values ± standard error of the mean (s.e.m) from at least 3 independent experiments performed; *p* values were derived from an unpaired two-tailed *t*-test, ****p-value<0.0001, **p-value<0.001, * p-value<0.05, n.s. no significant.

Then, we analyzed the expression levels of several ISC genes identified as IκBα targets in WT and IκBα KO HT-29 M6 cells at pre-confluence and 7 days of post-confluence. As expected, the majority of the ISC genes tested were down-regulated upon HT-29 M6 cell differentiation (induced by confluence). However, IκBα KO cells at 7-days post-confluence displayed ISC levels comparable to that detected in pre-confluent WT cells (Figure 5D and S5D). Further analysis of i-IκBα-repressed ISC genes indicated a general upregulation at 7-days post-confluence in IκBα KO compared with WT cells (Figure S5E).

We then studied whether specific IκBα activity affected *in vitro* tumor-initiating capacity of HT-29 cells. We seeded 600 single i-IκBα WT, SOF^ΔNF-κB^ or SOF^ΔH2A/H4^ cells in 3D cultures and maintained for 7 days in the absence or presence of doxycycline (to induce i-IκBα expression). Ectopic expression of i-IκBα WT significantly reduced the tumor-initiating capacity of IκBα KO cells, which was only marginally affected by expression of either SOF^ΔNF-κB^ or SOF^ΔH2A/H4^ (Figure 5E). i-IκBα SOF^ΔNF-κB^ but not SOF^ΔH2A/H4^ expression imposed a slight but significant reduction on the size of tumor spheres (Figure 5F) suggesting that chromatin-related IκBα function controls stemness/differentiation (leading to reduced proliferation) but both NF-κB and chromatin IκBα functions cooperate in regulating tumor-initiating capacity.

### *NFKBIA* deletion is associated with higher tumor initiation activity and poor patient survival in human cancer

To further confirm these results in a system closer to the patients, we moved to CRC patient-derived organoids (PDO). By IF analysis, we found that different PDO lines contain variable amounts of total and nuclear p-IκBα with PDO5 showing the highest nuclear p-IκBα levels (Figure 6A). Genetic deletion of IκBα by CRISPR-Cas9 in these cells (Figure 6B) led to transcriptional alterations (as determined by RNA-seq analysis of PDO5) signified by increased levels of canonical ISC markers such as *LGR5*, *PROM1* or *ASCL2* (Figure 6C). We then examined the *in vitro* tumor-initiating capacity of WT and IκBα KO PDO cells in Matrigel cultures. For all IκBα deficiency led to a significant increase in tumor-initiating capacity in al PDO lines tested, which was proportional to the number of cells seeded (Figure 6D), and maintained over passages (Figure S6A) indicative of long-term clonogenic activity. IκBα KO PDOs also displayed increased tumor size (Figure 6E) that was consistent with their higher proliferation rates (as determined by ki67 staining) (Figure S6B). We then examined a pan-cancer cohort encompassing 33 types of human cancer and found *NFKBIA* deletions in 2,244 (20.9%) of 10,746 tumors including 2,229 cases with hemizygous deletions and 15 cases with homozygous deletions (Figure 6F). *NFKBIA* deletions denoted a comparatively dismal prognosis in the 9,160 cases with available outcome data, in both a two-class and a continuous univariate Cox model (Figures 6G and 6H). Estimated median survival times for tumors with and without hemizygous *NFKBIA* deletions were 4.3 vs. 8.0 years, respectively. Hemizygous *NFKBIA* deletions remained independently associated with overall survival in adjusted Cox models that included cancer type, patient age, and gender, with or without pathologic stage as covariates (Figure 6H).

**Figure 6.**
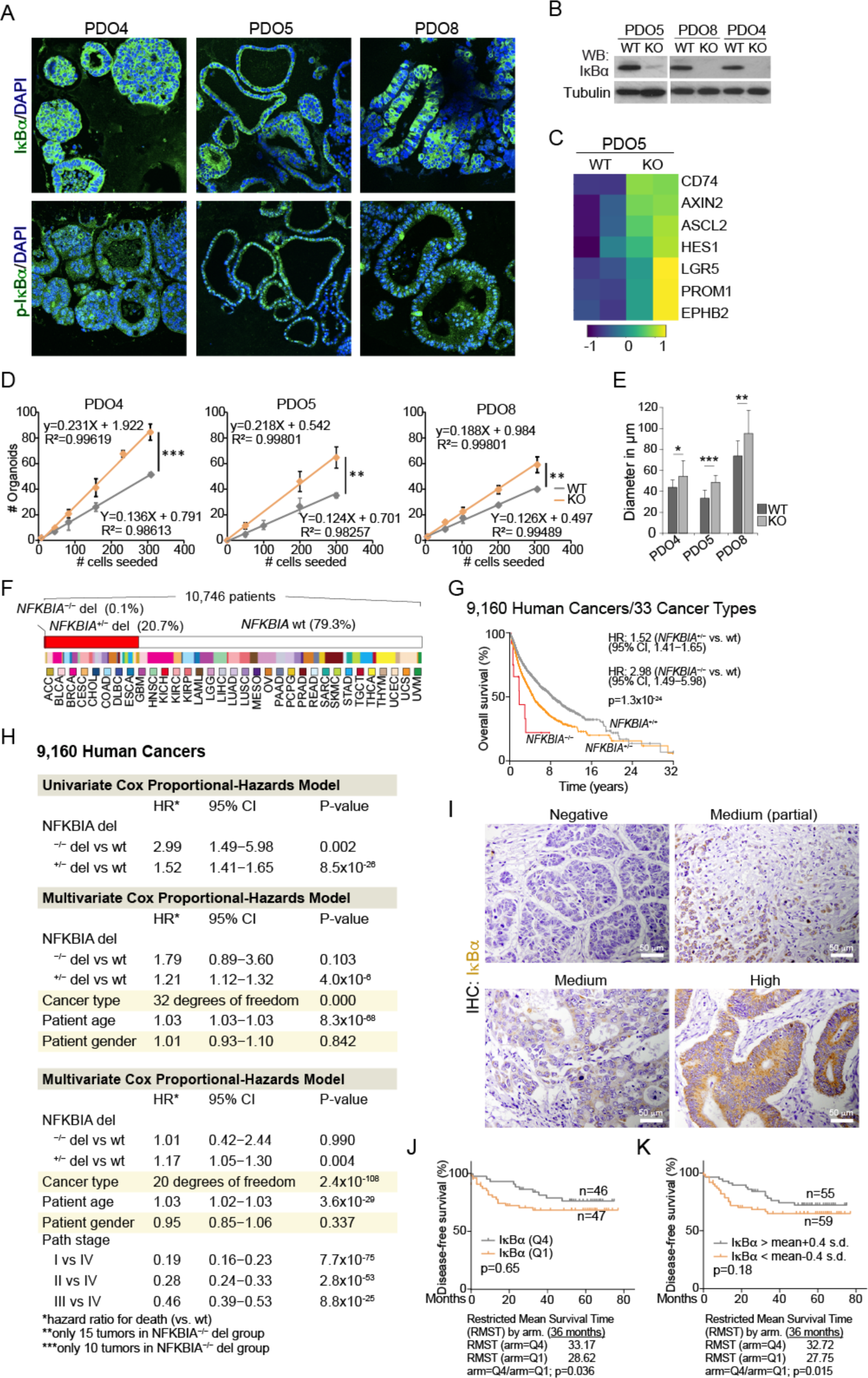
*NFKBIA* deletion is associated with higher tumor initiation activity and poor patient survival in human cancer. **(A)** IF analysis of total and p-IκBα in the indicated PDOs. **(B)** WB analysis of IκBα in WT and CRISPR-Cas9 IκBα-deleted PDO cells. tumors classified as low or high *NFKBIA*. **(C)** Heatmap representation of the expression levels of the indicated ISC genes in WT and IκBα-deleted PDO5 as determined by RNA-seq analysis. **(D, E)** Number (D) and diameter (E) of the organoids obtained by seeding the indicated number of WT and IκBα-deleted single PDO cells. **(F)** Incidence of *NFKBIA* deletions in a pan-cancer population. Note the high frequency of hemizygous deletions (+/-) when compared with homozygous deletions (-/-). ACC: adrenocortical carcinoma, CESC: Cervical squamous cell carcinoma and endocervical carcinoma, CHOL: Cholangiocarcinoma, COAD: Colon adenocarcinoma, DLBC: Diffuse Large B-cell Lymphoma, ESCA: Esophageal Carcinoma, GBM: Glioblastoma multiforme, HNSC: Head and Neck squamous cell carcinoma, KICH: Kidnet Chromophobe, KIRK: Kidney renal clear cell carcinoma, KIRP: Kidney renal papillary cell carcinoma, LAML: Acute Myeolid Leukemia, LGG: Brain Lower Grade Glioma, LIHC: Liver hepatocellular carcinoma, LUAD: Lung adenocarcinoma, LUSC: Lung squamous cell carcinoma, MESO: Mesothelioma, OV: Ovarian serous cystadenocarcinoma, PAAD: Pancreatic adenocarcinoma, PCPG: Pheochromocytoma and Paraganglioma, PRAD: Prostate adenocarcinoma, READ: Rectum adenocarcinoma, SARC: Sarcoma, SKCM: Skin Cutaneous Melanoma, STAD: Stomach adenocarcinoma, TGCT: Testicular Germ Cell Tumors, THCA, Thyroid carcinoma, THYM: Thymoma, UCEC: Uterine Corpus Endometrial Carcinoma, UCS: Uterine Carcinosarcoma, UVM: Uveal Melanoma. **(G)** Kaplan–Meier curves representing patient overall survival according to the presence/absence of homozygous (-/-) or hemizygous (+/-) *NFKBIA* deletions. P values were obtained using the Cox model likelihood-ratio test. **(H)** Univariate and adjusted Cox proportional-hazard models in 9,160 patients of the same population. Hazard ratios for death are displayed for the presence or absence of *NFKBIA* deletions. CI: Confidence Interval. The degree of freedom in the adjusted Cox model is restricted to 20 since pathological stage data was only available for 21 tumor types. **(I)** Representative images of CRC samples with the indicated IκBα staining patterns. **(J, K)** Kaplan–Meier curves representing patient disease-free survival according to the protein levels of IκBα as determined by multiplying the intensity of the staining (0-3) and the percent of the positive tumor area (0-100). We have either considered the quartile (J) and the average value plus/minus 0.4 or 0.6 standard deviations (K). In D and E, dots and bars represent mean values ± standard error of the mean (s.e.m) from at least 3 independent experiments performed; *p* values were derived from an unpaired two-tailed *t*-test, ***p-value<0.0005, **p-value<0.001, * p-value<0.05.

Finally, we performed IHC analysis in our *in-house* cohort of about 200 CRC samples with available clinical data to investigate the possibility that IκBα protein was a prognosis biomarker in CRC patients. We found variable levels and distribution of IκBα among tumor samples (Figure 6I) with low IκBα protein levels being sufficient to demarcate the subset of patients with poorest disease-free survival when comparing quartiles 1 and 4 (Figure 6J) or when considering the mean value +/- 0.4 standard deviations (Figure 6K). Prognosis value for the IκBα protein reached statistical significance when considering a 36-months follow up (Figures 6J and 6K, lower panels).

Collectively, we have here provided, for the first time, a conclusive demonstration that IκBα exerts a chromatin-related function on a subset of normal and cancer stem cell genes with impact in gene transcription, intestinal differentiation and tumor initiating activity. Importantly, we uncovered *NFKBIA* as a robust prognosis biomarker in a broad subset of human cancers.

## Discussion

For years, NF-κB inhibition has been considered the only IκBα function, which is shared by all other IκB homologs. However, IκBα deletion in mice imposes a striking phenotype that is not compensated by IκBβ, IκBε or p100 (IκBδ), suggesting that IκBα may exert additional NF-κB-unrelated functions. In support of this possibility, we previously demonstrated the presence of SUMOylated IκBα in the nuclear compartment of specific cell populations, including adult stem cells of the skin and intestine. Binding of SUMOylated IκBα to the chromatin takes place through direct association with histones H2A and H4 and provides cytokine responsiveness to a subset of PRC2 target genes, which are cell-type specific. For example, whereas SUMOylated IκBα regulates multiple genes of the *HOX* and *IRX* families as well as *NEUROG2* and *NEUROD4* in keratinocytes, it regulates *MEX3A* or *BMI1* in intestinal cells, thus controlling tissue stem cell maturation and lineage cell differentiation^9,11^. Further supporting the concept that IκBα exerts both NF-κB-related and -unrelated functions, two IκBα homologs are present and functional in the nematode *C.elegans*^10^, which lacks recognizable NF-κB factors (see https://www.bu.edu/nf-kb). Moreover, the homeotic phenotype impose by Cactus (IκBα) hemizygous deletion in Drosophila is not rescued by reducing dorsal (p65-NF-κB) levels^9^.

In pathology, loss of nuclear IκBα and accumulation of cytoplasmic IκBα correlates with cancer progression in the human skin and it is linked to aberrant *HOX* gene transcription^9^, which has been canonical targets of PRC2 activity. However, the prominent function of NF-κB in gene regulation and the multiple cross-talks between NF-κB and polycomb^12, 14^ have precluded, until now, the study of NF-κB-independent IκBα functions (i.e., PRC2 related) not only in cancer but also in the different physiological systems where it plays a role. We have now performed a sophisticated bioinformatic analysis of the evolutionary conserved residues of the IκBα protein to uncover the most likely contributors to relevant protein-protein interactions. This analysis indicated that IκBα binds to NF-κB and histones through the same region, which we had already suggested based on its mutually exclusive binding to either partner^9^. Strikingly, residues essential for NF-κB binding were conserved during evolution and among the different IκB homologs including BCL3. However, residues involved in histone binding were exclusively found in the IκBα homolog but are still evolutionary conserved, indicating that the chromatin function of IκBα represents an exclusive but ancestral NF-κB-independent function.

Using separation-of-function IκBα mutants, we have shown that goblet cell differentiation requires the chromatin function of IκBα (rescued by IκBα WT and the SOF^ΔNF-κB^ mutant), which allows the proper transcriptional repression of ISC genes. However, transcriptional ISC repression imposed by IκBα is previous to the detection of H3K27me3 /the mark deposited by PRC2) indicating that additional chromatin-related functions of IκBα (i.e. recruitment of histone deacetylases or other chromatin-editing enzymes) may contribute to gene regulation.

We here establish that IκBα SOF mutants represent a unique tool for basic research (i.e., in the stem cell and development field) and clinical research, including the more specific cancer models where IκBα deletions are highly prevalent such as Hodgkin lymphoma^33, 34^ or glioblastoma^16, 35^ and also neuronal disorders where defective IκBα function plays a role. The design of a new tool to reconstitute either NF-κB or PRC2 regulation in IκBα-depleted cancers will allow a diagnosis refinement (i.e., stratification of tumors fueled by aberrant NF-κB or PRC2 activity) and a better therapeutic prescription for specific groups of patients.

## Author contributions

L.E. and A.B. conceptualized the study, designed the experiments and wrote the manuscript. M.F and J.V. performed bioinformatic analysis to identify IκBα interaction domains. D.A-V., J.B., L.S., L.G-P. and L.M. generated the IκBα mutants and performed *in vitro* experiments and functional analysis. T.L-J and M.M. analyzed transcriptomic data. M.B. and R.R. contributed to the experimental design, data interpretation and manuscript writing. D.A-V., L.S., T.L-J., L.G-P. and L.E. prepared the figures.

## Acknowledgments

We want to thank Espinosa’s and Bigas’ lab members for constructive discussions and suggestions, and Nuria Mascaró for technical support. This work was funded by grants from Instituto de Salud Carlos III FEDER (PI19/0013), Generalitat de Catalunya 2017SGR135 and CIBERONC. D.A-V is a recipient of the FI20/00130 grant from Instituto de Salud Carlos III FEDER. T.L-J is a recipient of the AECC postdoctoral grant POSTD21975. L.S. is a postdoctoral researcher from CIBERONC.

## MATERIALS AND METHODS

### Cell lines and reagents

All cells were grown in Dulbecco’s modified Eagle’s medium (DMEM) [Invitrogen] supplemented with 10% fetal bovine serum (FBS) [Biological Industries]. Cells were grown in an incubator at 37°C and 5% CO2. Cells used in these studies were HEK-293T [ATCC Ref. CRL-3216], HCT-116 [ATCC Ref. CCL-247] and HT-29 M6 cells derived from parental HT-29 [ATCC Ref. HTB-38D] by adaptation to methotrexate^36^.

### Generation of SOF^ΔNF-κB^ and SOF^ΔH2A/H4^ IκBα mutants and stable inducible cell lines

Human IκBα cDNA was cloned into pPiggybacTRE vector^37^ as a XbaI/SalI fragment into NheI-SalI sites, using standard procedures. The XbaI and SalI sites were incorporated at the ends of the cDNA as 5’ extensions in PCR primers (Table S4). To generate the SOF mutants, we used sequence overlap extension followed by PCR using the mutagenic primers shown in Table S4. IκBα KO HT-29 M6 cells^27^ were reconstituted using the piggybac constructs (encoding either inducible (i-)IκBα WT or one of the SOF mutants) along with the transposase to increase integration efficiency. Briefly, cells were transfected with a mixture 1:1 of the transposase coding plasmid and the corresponding piggybac plasmid. Two days after transfection, hygromycin (400 µg/mL) was added and cells were cultured for three weeks with several medium changes to eliminate dead cells and refresh hygromycin. Clones were plucked, expanded and screened for i-IκBα expression.

Transfections were performed using Polyethylenimine (PEI) [Polysciences Inc. Ref. 23996] following standard procedures. Briefly, we mixed 4μg of PEI (as a 1mg/mL suspension in water) per μg of DNA in serum-free DMEM and incubated 5 min at room temperature. Then, The PEI dilution was added to DNA, previously placed in a fresh tube (1 µg/well for 6 well plates or equivalent DNA mass/culture surface), and incubated for 20 min. at RT. Finally, we added the PEI/DNA mix to cells.

### IκBα knock-in cell lines

To modify the *NFKBIA* gene *in situ*, two guide RNAs were designed using the Benchling platform (https://www.benchling.com/crispr/). One guide targets the double DNA cut to 39 nucleotides 3’ of the splice donor for intron 2 and the other to 4 nucleotides 3’ of the ATG codon in exon 1 in the original gene (NC_000014.9). The sequence to produce these guide RNAs, primers Sg_1*NFKBIA* and Sg_2*NFKBIA* respectively (Table S4), was introduced in the pLentiCRISPR V2 plasmid following published procedures^38^. In addition, 4 repair plasmids, pBS-*NFKBIA*-ki WT, N108A, M3 and β-mimic were constructed to achieve homologous recombination next to the targeted chromosomal locations.

These plasmids carry two homology arms which extend 1043 bp upstream and 1043 bp downstream of the fragment to be replaced. Between the two homology arms there is: i) a loxp site followed by the 8 nucleotides that precede the ATG codon, the second *NFKBIA* codon TTC and the GFP coding sequences; ii) peptide 2A coding triplets in frame with the GFP coding codons followed by a second loxp site; and iii) *NFKBIA* gene sequences starting with the 8 nucleotides that precede the ATG triplet and continuing till the cutting site determined by the downstream guide RNA used. A deletion of two intronic cytosines destroys the targeted PAM in the complementary strand. The sequence corresponding to triplets encoding amino acids 84 to 108, laying between the two homology arms, has been tailored to yield the desired mutant protein in edited cells.

### CRISPR/Cas9-mediated *NFKBIA* gene edition

HCT116 cells were co-transfected with the described targeting plasmids and the appropriate repair plasmid at a 1:1:40 molecular ratio. After 24 hours incubation, puromycin (2 μg/mL) was added for 72 hours to enrich for transfected cells and then removed. Cells were expanded for about two weeks, sorted for GFP expression, and plated as single cells in 96-well plates to isolate clones. Clones were genotyped by selective amplification and Sanger sequencing of chromosomal fragments that expand beyond the homology arms in repair plasmids. Primers used to construct two pLentiCRISPR V2 derivatives able to provide transfected cells with single guides (Sg) for genetic edition of the *NFKBIA* gene are specified in Table S4.

### Pull down and peptide-based immunoprecipitation (IP)

PD assays were performed as previously described^39^. Briefly, GST fusion proteins were incubated with lysates for 45 min in a rotary shaker at 4°C. When indicated, nuclear extracts were boiled at 98°C for 5 min in the presence of 1% SDS to disassemble pre-existing protein complexes and then neutralized in 1% Triton X-100. Precipitates were resolved in SDS-PAGE and analyzed by IB. For peptide IP, histone H4 peptides were synthesized as biotinylated N-terminal and C-terminal amides. Peptides were incubated overnight at 4**°**C with the indicated cell extracts and precipitated with streptavidin-Sepharose beads for 45 min.

### Cell fractionation and Western Blot (WB)

For soluble and chromatin separations, cells were lysed 1mM EDTA, 0.1mM Na-orthovanadate (Na3VO4), 0.5% Triton X-100, 20mM β-glycerol-phosphate, 0.2mM PMSF, protease inhibitor cocktail, in PBS for 20 min on ice and centrifuged at 13,000 rpm. Supernatants were recovered as the soluble fraction, and the pellets were lysed in Laemmli buffer (1x SDS-PAGE buffer plus β-mercaptoethanol (BME) [Sigma, Ref. M-3148]) or in 1% SDS PBS, sonicated and treated with 1% Triton X-100. Lysates were analyzed by Western blotting using standard SDS–polyacrylamide gel electrophoresis (SDS-PAGE) techniques. In brief, protein samples were boiled in Laemmli buffer, run in polyacrylamide gels, and transferred onto polyvinylidene-difluoride (PVDF) membranes [Millipore Ref. IPVH00010]. Membranes were incubated overnight at 4°C with the appropriate primary antibodies, extensively washed and then incubated with specific secondary horseradish peroxidase–linked antibodies from Dako [Ref. P0260 and P0448]. Peroxidase activity was visualized using the enhanced chemiluminescence reagent [Biological Industries Ref. 20-500-120] and autoradiography films [GE Healthcare Ref. 28906835].

### Immunofluorescence (IF) analysis

Tissues were fixed in 4% formaldehyde overnight at room temperature and embedded in paraffin. 4μm paraffin embedded sections were first de-paraffinized in xylene. IHC was performed following standard techniques with EDTA- or citrate-based antigen retrieval and developed with the Envision+ System HRP Labelled Polymer anti-Rabbit [Dako Ref. K4003] or anti-Mouse [Dako Ref. K4001] and developed with TSA^TM^ Plus Cyanine 3/Fluorescein System [PerkinElmer Ref. NEL753001KT] and mounted in ProLong^TM^ Diamond Antifade Mountant plus DAPI [Thermo Scientific Ref. P36971]. Images were taken in an SP5 upright confocal microscope (Leica).

For organoid direct immuno-fluorescence, we followed a modified protocol from Dow and colleagues ^40^. In brief, organoids were fixed with 4% paraformaldehyde, incubated with DTT buffer (100mM Tris pH9.4, 10mM DTT in H_2_O), permeabilized with 0.5% Triton X-100 [Thermo Scientific, Ref. 28340], washed and incubated overnight with the corresponding primary antibodies. As secondary antibodies we used Alexa Fluor^TM^ [Molecular probes, Ref.A21202 and A21206]. ProLong^TM^ Diamond Antifade Mountant plus DAPI [Thermo Scientific Ref. P36971] was used as mounting medium. Images were taken in an SP5 upright confocal microscope (Leica).

### Antibodies

Antibodies used were anti-IκBα (sc-371), anti-IκBε (sc-7275), anti-p50 (sc-7178), anti-p65 (sc-109) and anti-Lamin B (sc-6216) from Santa Cruz; anti-p-IκBα (#9246) from Cell Signaling; anti-HA (MMS-101R) from Babco; anti-SUZ12 (ab12073) from Abcam and (C15310029) from Diagenode, anti-Tubulin (clone B-5-1-2) from Sigma Aldrich and anti-MUC5AC (clone 45M1) from Neomarkers, Waltham, MA.

### RT-qPCR analysis

Total RNA from the cells of interest was extracted with the RNeasy Micro Kit, and cDNA was produced with the RT-First Strand cDNA Synthesis Kit. RT-qPCR was performed in LightCycler 480 system using SYBR Green I Master Kit. Samples were normalized to the mean of the housekeeping genes *TBP*, *HPRT1 and/or GAPDH*. Primers used for RT-qPCR and ChIP-PCR analysis are listed in Supplementary Table S5.

### RNA-seq experiments

Total RNA from untreated and doxycycline-treated i-IκBα WT, i-IκBα SOF^ΔH2A/H4^ and i-IκBα SOF^PCR2^ HT-29 M6 IκBα KO cells was extracted using RNeasy Micro Kit. The RNA concentration and integrity were determined using Agilent Bioanalyzer [Agilent Technologies]. Libraries were prepared at the Genomics Unit of PRBB (Barcelona, Spain) using standard protocols and cDNA was sequenced using Illumina HiSeq platform (75 bp paired-end reads). Samples sequencing depth ranged between 36.1M and 47.9M reads (average 44.1M reads) per sample.

Quality control was performed on raw data with FASTQC tool (v0.11.9). Raw reads were trimmed to remove adapters presence with TrimGalore (v0.6.6)^41^. Default parameters were used except for the required minimum quality and the stringency parameters which were set to 15 quality score and 3 bases respectively. Trimmed reads were aligned to reference genome with STAR aligner tool (v2.7.8)^42^. STAR was executed with default parameters except for the number of allowed mismatches which was set to 1. Required genome index was built with corresponding GRCh38 gtf and fasta files retrieved from Ensembl (http://ftp.ensembl.org/pub/release-106/). Obtained BAM files with uniquely mapped reads were considered for further analysis. Raw gene expression was quantified using featureCounts tool from subRead software (v2.0.1)^43^. Those genes with less than 10 counts among all samples were removed for downstream analysis.

Differential expression analysis was conducted with DESeq2 R package (v1.34)^44^. Comparisons between treatment conditions per cell type were considered. For visualization purposes, counts matrix was normalized with the Variance Stabilizing Transformation. Plots were done in R. Expression heatmaps were generated using the heatmaply package in R (v. 1.3.0)^45^. GSEA was performed with described ISC gene set using gene set permutations (n=1000) for the assessment of significance and log2_ratio_of_classes metric for ranking genes. RNA-sequencing data for i-IκBα WT and SOF mutant HT-29 M6 cells are deposited at the GEO database with accession number GSE206515 (token: ilyrmckidlyjjax).

### ChIP-seq experiments

ChIP for H3K27me3 was performed as previously described^9,11^. Briefly, formaldehyde crosslinked cell extracts were sonicated, and chromatin fractions were incubated for 16 hours with anti-H3K27me3 antibody [Millipore 07-449] in RIPA buffer and then precipitated with protein A/G Sepharose [GE Healthcare, Refs. 17-0618-01 and 17-0780-01]. Cross-linkage was reversed, and 6–10 ng of precipitated chromatin was directly sequenced in the genomics facility of Parc de Recerca Biomedica de Barcelona (PRBB) using Illumina Hi-Seq platform. Raw single-end 50-bp sequences were filtered by quality (Q > 30) and length (length > 20 bp) with Trim Galore^46^. Filtered sequences were aligned against the reference genome (mm10 release) with Bowtie2^47^. MACS2 software^48^ was run first foreach replicate, and then combining all replicates, using unique alignments (q-value < 0.1). Peak annotation was performed with ChIP seeker package^49^, and functional enrichment analysis with enrichR^50^, using the latest version of GO annotations. ChIP sequencing data are deposited at the GEO database with accession number GSE206515.

### Weighted contact number

The weighted contact number (WCN) is a metric that quantifies the contact density for a particular atom or residue in a protein structure. It is defined as the sum of all inverse squared distances of all other particles in the system to that specific atom:

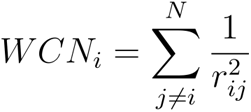

Here, *N* is the number of atoms in the structure and *r_ij_* is the distance between atoms *i* and *j*. The profiles are then normalized by calculating their standard score (z-score). Since we are only interested in making comparisons at the residue level, we only considered the alpha carbon atoms of the protein structures.

The WCN metric, as described here, has an inverse relationship with the evolutionary sequence conservation^40^ therefore, we use the inverse of the WCN (_i_WCN) to make direct comparisons to the evolutionary information.

### Sequence Conservation score

The sequence conservation scores were derived from the CONSURF server.^41^ First, the method builds phylogenetic trees from multiple sequence alignment of homologous sequences to the target protein. Then, and considering the stochasticity underlying the evolutionary process, conservation profiles are derived using the Empirical Bayesian Method and smoothed over a window of five residues^42^. Finally, the scores are normalized to their corresponding z-scores; thus, the Sequence Conservation score for residue *i*(*SC_i_*) corresponds to the z-score value calculated from the distribution of CONSURF scores.

### Fold-Excluded Evolutionary Conservation (FEEC) Score

Residues conserved in a particular set of homologous proteins need to fulfill a relevant thermodynamic or kinetic role for the given biomolecular system. This role can be the maintenance of the folded structure, or it can also regulate other phenomena such as binding, catalysis, solubility, folding kinetics, etc. A three-dimensional protein structure contains the chemical contact information necessary to adopt its particular fold. The greater the number of chemical contacts a residue participates in, the more restricted it is to change its identity without compromising essential interactions that support the folded structure. Likewise, if the residue is not involved in many chemical interactions, it can vary more freely along the structurally-permitted sequence landscape.

The atomic contact density, measured here as the _i_WCN metric, has been shown to correlate well with dynamic and sequence conservation of proteins^40, 43^. The later correlation is higher when the _i_WCN values are derived using the full biological complex structure^40^. Since the _i_WCN only considers the contact information derived from a particular protein structure, residue positions with higher sequence conservations than the _i_WCN metric should have different or additional roles than maintaining the folded configuration (e.g., intermolecular binding). Based on this logic, we defined a Fold-Excluded Evolutionary Conservation (FEEC) score as the value of the _i_WCN minus the Sequence Conservation (SC) score: *FEEC_i_* = *iWCN_i_* − *SC_i_*

Based on this definition, if a residue’s FEEC score is positive, it could have additional conservation constraints than those imposed by the protein’s tertiary structure alone.

### Analysis of residue interface score

We obtained a set of interacting conformations between IκBα and NF-κB to define an interacting energy landscape. For this, we generated 6500 minimization trajectories from the crystallographic complex (PDB code: 1NFI) using the relax protocol of the Rosetta software.45. We then assign a probability to each complex conformation from a Boltzmann distribution based on the entire energy landscape of the ensemble of sampled conformations:

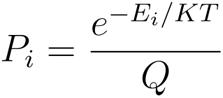

where *E_i_* is the energy of the i^th^ conformation, *KT* is the energy partition constant (set equal to one), and *Q* is the partition function of the respective ensemble of N structures:

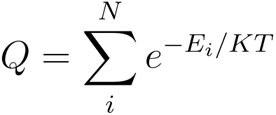

The interface binding energy 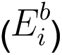 for each complex structure is calculated as the difference between the energies of the interacting complex structure 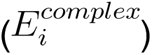 and the individual protein energies 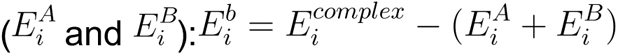

All interface energies were then integrated using the complete set of probabilities to calculate the binding energy expectation value (interface score):

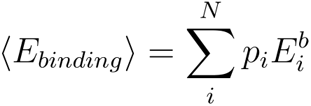

We employed the Rosetta energy function^44^ in all the modeling steps. This function is residue decomposable, and therefore it is straightforward to do the energy analyses only considering the contribution of individual residues. Accordingly, per-residue interface scores were obtained as the expectation values of individual residues’ binding energies.

### In-silico analysis of the effect of mutating non-synonymous residues of IκBα in its structure

The protocol includes the minimization of the WT structure and the generation of selected mutants by changing the identity of the target residue. This is followed by repacking and backbone minimization of the target and neighboring residues until a radius of 8 angstroms. The energy differences between mutating to the WT amino acid and to any of the other 20 amino acids are reported.

## Competing Interests

Laura Solé, Anna Bigas and Lluís Espinosa have a pending patent application related with this work (EP2023/055330). The authors have no additional financial interests.

## Data and Materials Availability

All data associated with this study are present in the paper or the Supplementary Materials. RNA sequencing and ChIP sequencing data have been deposited in NCBI’s Gene Expression and are accessible through GEO Series accession no. GSE206515.

## SUPLEMENTARY FIGURES AND LEGENDS

**Figure S1.**
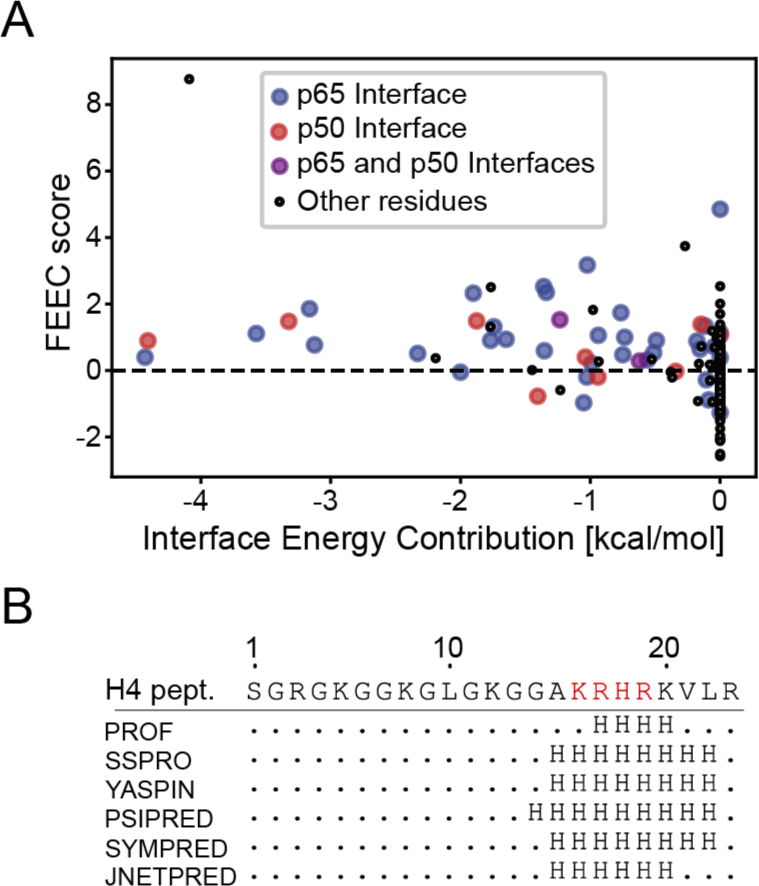
Structural characterization of the N-terminal tail of histone H4. **(A)** Classification of IκBα and NF-κB interface residues based on the Fold-excluded Evolutionary Conservation (FEEC) score. All IκBα residue energy contributions to the interface with NF-κB are plotted against their respective FEEC scores. Interface residues between IκBα and NF-κB subunits appear in blue (p65 subunit), red (p50 subunit), or purple (p65 and p50 subunits), all remaining residues appear in black circles. **(B)** Secondary structure prediction of the N-terminal peptide of the H4 Histone (H4p). Positions indicated with an H have a helical character according to the prediction method. The NLS motif is displayed as red letters in the H4p sequence.

**Figure S2.**
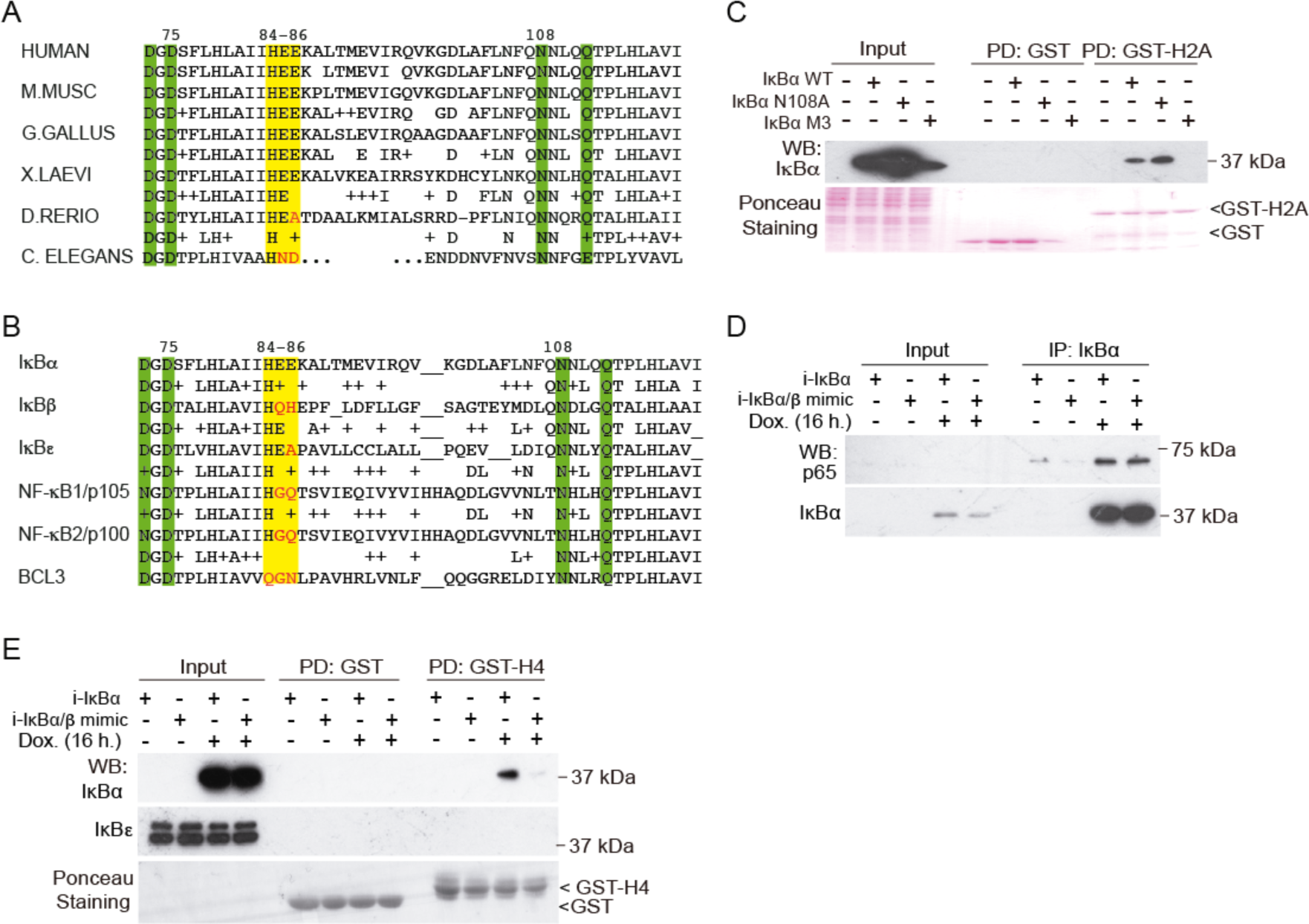
Two different evolutionary conserved Aa clusters of IκBα define NF-κB or histone H2A and H4 binding. **(A)** Sequence alignment of the IκBα sequence (Aa. 73 to 121 in the human orthologue) in the indicated species. Notice the absolute conservation of residues D75 and N108 (essential for NF-κB binding) and the cluster formed by the acidic residues HEE (essential for histone H4 and PRC2 binding). **(B)** Sequence alignment of the IκBα sequence (Aa. 73 to 121 in the human orthologue) with the indicated IκB homologs. **(C)** PD assay using GST, GST-H2A as bait and the indicated i-IκBα SOF cell lines. We determined the presence of IκBα in the precipitates by WB. **(D, E)** Co-IP using the anti-IκBα antibody (D) and PD assay using GST-H4 as bait (E) to determine the relative binding capacity of the IκBα (β-mimic) to p65 (NF-κB) and histones. Notice the high reduction in the binding capacity of the mutant to histone H4 when compared with WT IκBα. Ponceau staining in C and E are shown to demonstrate comparable amount of GST and GST-H2A/GST-H4 in the assays.

**Figure S3.**
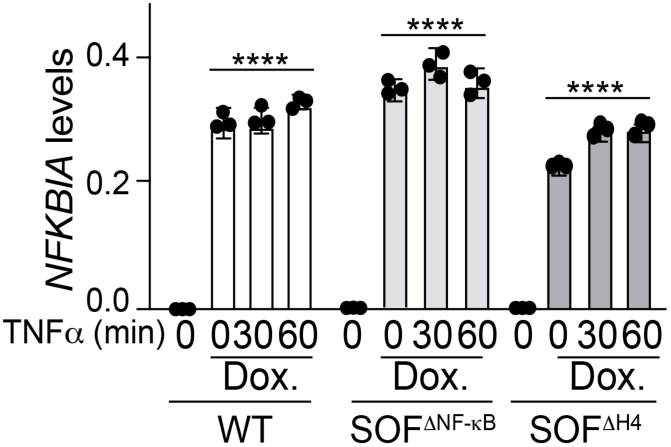
Similar transcriptional induction of IκBα WT and SOF mutants after doxycycline treatment. qPCR analysis of IκBα expression (*NFKBIA* gene) in IκBα KO HT-29 M6 cells reconstituted with i-IκBα WT or the indicated SOF mutants, left untreated or treated for 16 hours with doxycycline (Dox.). *NFKBIA* levels were calculated relative to *GAPDH*. Bars represent mean values ± standard error of the mean (s.e.m) from at least 3 independent experiments performed; *p* values were derived from an unpaired two-tailed *t*-test, ****p-value<0.0001.

**Figure S4.**
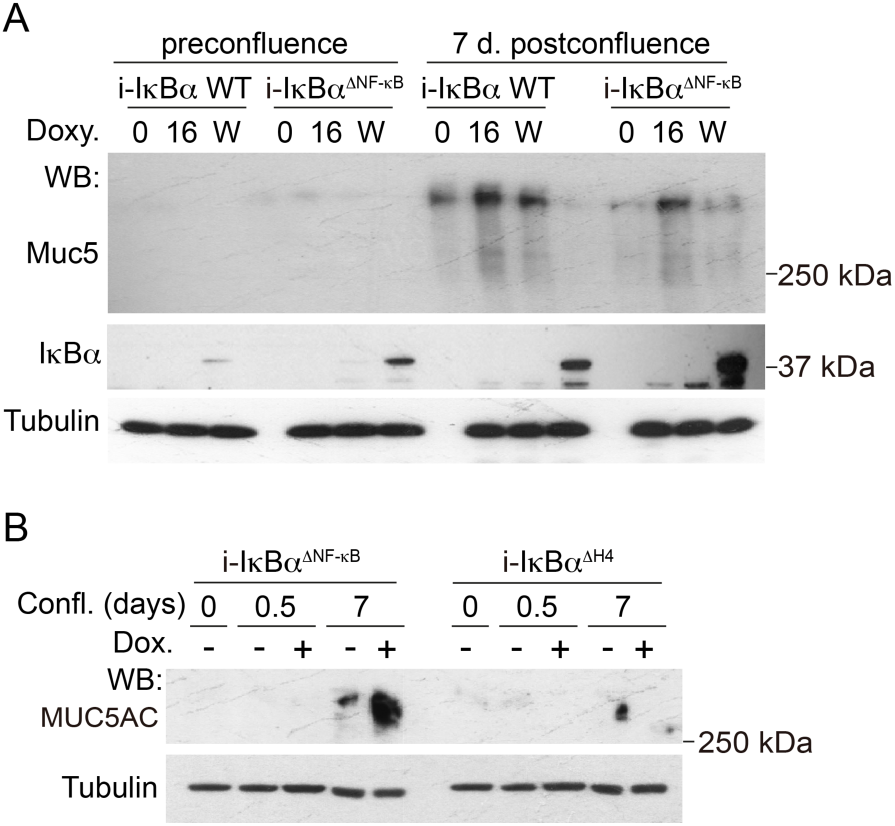
i-IκBα SOF^ΔNF-κB^ mutant rescues defective differentiation imposed by IκBα deficiency. **(A, B)** Western blot analysis of the differentiation marker MUC5AC in HT-29 M6 IκBα KO cells reconstituted as indicated and collected at pre-confluence or at 0.5 and 7 days of post-confluence. In A, i-IκBα expression was induced by treating cells with doxycycline for 16 hours (16), for the whole period of the experiment (W), or left uninduced (0). In B, doxycycline was added for 16 hours at pre-confluence.

**Figure S5.**
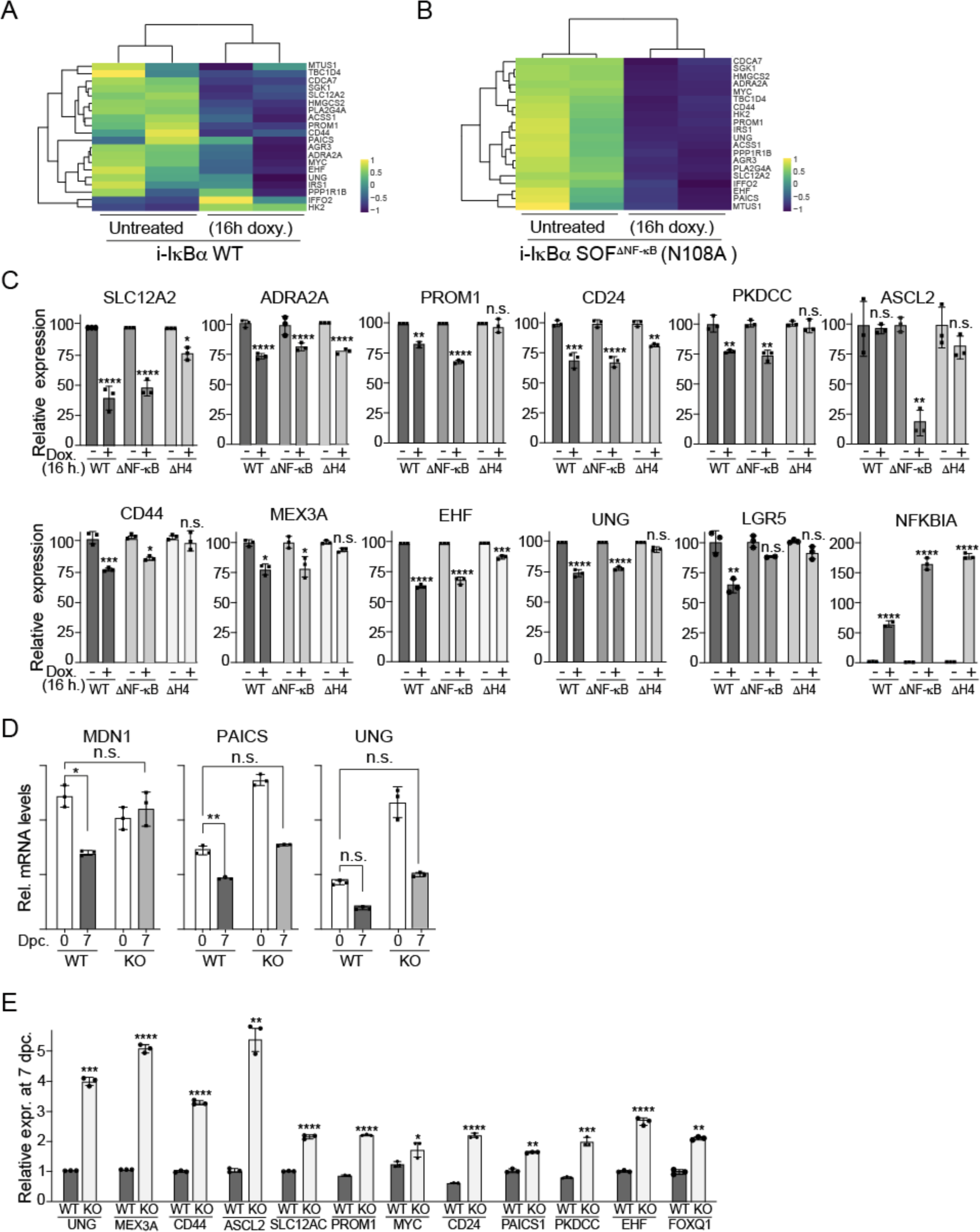
Differential regulation of ISC genes upon IκBα modulation of human intestinal cancer °cells. **(A, B)** Heatmap of the 20 top ISC genes that were downregulated in KO cells reconstituted with i-IκBα SOF^ΔNF-κB^ (A). Heatmap of the same 20 genes in the i-IκBα WT expressing cells (B). **(C)** qPCR of IκBα-regulated ISC genes (identified in the RNA-seq analysis) in IκBα KO HT-29 M6 cells reconstituted with the indicated i-IκBα species. **(D, E**) qPCR analysis of IκBα-regulated ISC genes in WT and IκBα KO HT-29 M6 collected at pre-confluence (0) and 7 dpc (D) or at 7 days of post-confluence (7 dpc) (E). In C-E, bars represent mean values ± standard error of the mean (s.e.m) from at least 3 independent experiments performed; *p* values were derived from an unpaired two-tailed *t*-test, ****p-value<0.0001, ***p-value<0.0005, **p-value<0.001, * p-value<0.05, n.s. non-significant.

**Figure S6.**
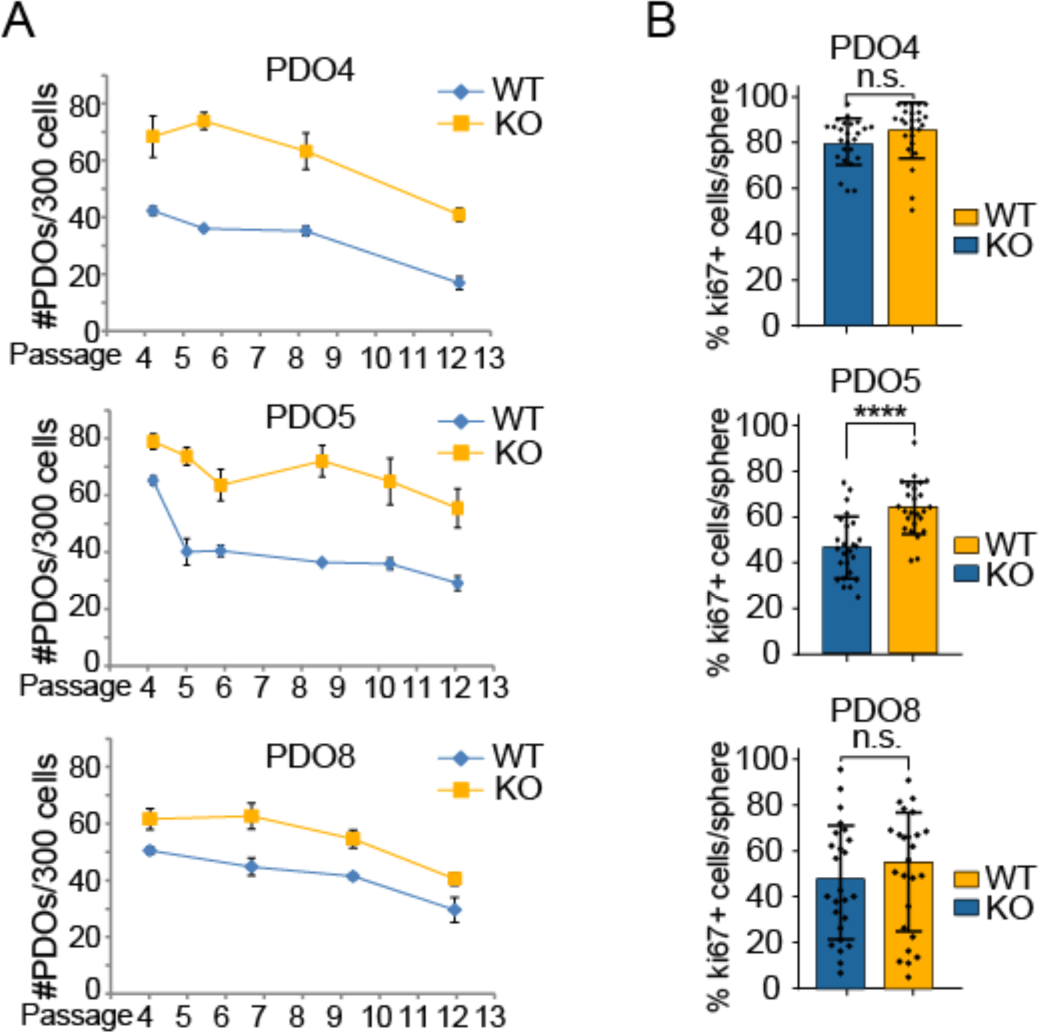
Effect of *NFKBIA* deletion in tumorigenic activity. **(A)** Clonogenic assay of the indicated PDO cells after serial single-cell passaging. **(B)** Quantification of the percentage of proliferating cells determined by Ki67 staining in PDO cells. P-value was calculated by unpaired two-tailed *t*-test. n.s., non-significant; ****, p<0.0001.

## Notes

### Competing Interest Statement

Anna Bigas, Laura Sole and Lluis Espinosa have a submitted patent related with SOF IκBα mutants

