## Supplemental Table S4 for "Separation-of-function mutants reveal the NF-κB-independent involvement of IκBα in the regulation of stem cell and oncogenic programs"

**Table 3: Primers used to generate inducible IκBα expression plasmids and knockin constructs**

| *NFKBIA* FW XbaI | aatataTCTAGAgccaccatgttccaggcggccgagcg |
| --- | --- |
| *NFKBIA* RV SalI | aatataGTCGACtaacgtcagacgctggc |
| *IKBA* 84-86A_FW | ttcctgcacttggccatcatcGCtgCagCaaaggcactgaccatggaa |
| *IKBA* 84-86A_RV | ttcctgcacttggccatcatcGCtgCagCaaaggcactgaccatggaa |
| *IKBA* 108A_FW | GCCTTCCTCAACTTCCAGgcCAACCTGCAGCAGACTCC |
| *IKBA* 108A_RV | GGAGTCTGCTGCAGGTTGgcCTGGAAGTTGAGGAAGGC |
| *IKBA* 72A_FW |  |
| *IKBA* 72A_RV |  |
| *IKBA* 73A_FW |  |
| *IKBA* 73A_RV |  |
| *IKBA* 75A_FW |  |
| *IKBA* 75A_RV |  |
| *IKBA* β-mimic_FW | gcacttggccatcatccatCaaCaTaaggcactgaccatggaag |
| *IKBA* β-mimic_RV | cttccatggtcagtgccttAtGttGatggatgatggccaagtgc |
| Sg_1*NFKBIA* _FW | CACCGgctacatcagctacgtccca |
| Sg_1*NFKBIA* _RV | AAACtgggacgtagctgatgtagcC |
| Sg_2*NFKBIA* _FW | CACCGcgtccgcgccatgttccagg |
| Sg_2*NFKBIA* _RV | AAACcctggaacatggcgcggacgC |
