## Supplemental Table S5 for "Separation-of-function mutants reveal the NF-κB-independent involvement of IκBα in the regulation of stem cell and oncogenic programs"

**Table S4. Primers used in the qPCR analysis**

| **Target Gene** | **Forward** | **Reverse** |
| --- | --- | --- |
| **MEX3A** | TGAATCCTCCATGCAAATCA | GGTGCAAGGCCTGTAACAAT |
| **NFKBIA** | AAATACCCCCCTACACCTTGCC | CATCAGCCCCACACTTCAACAG |
| **CD24** | AACTAATGCCACCACCAAGG | CCTGTTTTTCCTTGCCACAT |
| **FOXQ1** | TGTGGCATTTCCAGGTATGA | GCCCAAGGAGACCACAGTTA |
| **PKDCC** | GGCAGCTGGTCTTTTTCAAG | TGGCATTATTGCACGTTTGT |
| **PODXL** | AAAGGCCAAAAGCTCAGACA | GGACGTTCCCACAACAGTCT |
| **CD44** | TGAATATAACCTGCCGCTTTG | CCGTCCGAGAGATGCTGTAG |
| **LGR5** | ATGGTCGCTCTCATCTTGCTC | ATATTCTCCAGGTCTCCCTTGTC |
| **A20** | GAGAGGCGGCAAAAGAATCAAAAC | TGAACAGAAAAGGGCTGGGTGC |
| **EHF** | TGCAGCATCTGAAGTGGAAC | AGGAAGGTACTGGTGGTTG |
| **UNG** | AATGGCAGCTGTATCCAACC | CACCCCAACATCTGTCACTG |
| **SLC12A2** | TGGTGGTGCAATTGGTCTAA | TTTTGCTTCCCACTCCATTC |
| **ADRA2A** | ACTGGACTACAAGGGCATGG | ACATCAAAACCAAGGCCAAG |
| **PROM1** | GCCAGCCTCAGACAGAAAAC | CCAAGCCTTAGGAGCATCTG |
| **ASCL2** | AGCTGGTGAACTTGGGCTTC | CTCCACCTTGCTCAGCTTCTT |
| **MYC** | AGCGACTCTGAGGAGGAACA | CCCTCTTGGCAGCAGGATAG |
| **MDN1** | CGTCTGACTCCTCAGGGAAG | AAGCATAAACCGAGGGTGTG |
| **PAICS1** | AGGATAATGGCGACAGCTGAG | GGCAGTTTTAATACCTGCTTCCT |
| **GAPDH** | GTCATCCCTGAGCTGAACG | CTCCTTGGAGGCCATGTG |
| **TBP** | TGCCCGAAACGCCGAATATAATC | GTCTGGACTGTTCTTCACTCTTGG |
